# Nonlinear rheology of laminin-111-modified hyaluronan hydrogels

**DOI:** 10.64898/2026.06.02.729677

**Authors:** Jordan L. Shivers, Mary C. Farach-Carson, Fred C. MacKintosh, Danielle Wu

## Abstract

We experimentally assess the nonlinear rheology of composite biopolymer hydrogels composed of thiolated hyaluronic acid, poly(ethylene glycol) diacrylate (PEGDA), and laminin-111 in varied concentrations. We focus in particular on the influence of laminin on the mechanics of the assembled hydrogels, reporting nonlinear rheological measurements for gels under applied shear and compressive load. We find that increasing the concentration of laminin in the synthesized gels reduces the linear shear modulus and gives rise to a mild strain softening regime at intermediate strains prior to the onset of strain stiffening. In the stiffening regime, we find that all gels exhibit stress-controlled mechanics with *K* ∝ *σ*^*a*^, with an apparent stiffening exponent of *a* ≈ 1, in agreement with observations of a variety of other reconstituted biopolymer gels. We discuss the possible implications of this nonlinear mechanical behavior on mechanotransduction and organoid development in biomimetic extracellular matrices.

## I. INTRODUCTION

The extracellular matrix (ECM) is a composite network consisting of several types of biopolymers with distinct mechanical properties. The mechanical properties of composite materials generally differ considerably from those of isolated components. Interactions between components often give rise to qualitatively unexpected behavior, such as enhanced toughness and synergistic stiffening [2, 3] in multicomponent hydrogels. Understanding the individual and cooperative mechanical contributions of the various ECM components can bolster efforts to design artificial, biomimetic ECMs for applications in tissue engineering, studies of cell-matrix interactions, and uses for organoid growth and differentiation. The mechanical response of biological matrices under large, nonlinear deformations is especially important in the context of the growth and morphogenesis of embedded cells or organoids [4], in which active shape changes can strongly deform the surrounding matrix [5]. While the mechanics of type I collagen hydrogels have been well characterized in the large-strain nonlinear regime [6], comparatively little is known about the nonlinear mechanics of other components commonly used in reconstituted ECM models. A commonly used ingredient in biomimetic ECMs is hyaluronic acid or hyaluronan (HA), shown schematically in Fig. 1, a flexible polyelectrolyte that imbues living tissues with resistance to compression and strongly influences their development and morphogenesis [7–21]. Often, HA used in the context of artificial ECMs is crosslinked by active agents such as poly(ethylene glycol) diacrylate (PEGDA) [22, 23], sketched in Fig. 1. Laminin-111 (which we will refer to as laminin), also sketched in Fig. 1, is a trimeric glycoprotein that plays a crucial role in mediating cell-matrix interactions in the basement membrane [7, 24]. When isolated, laminin polymerizes into weakly connected networks that do not alone form a hydrogel [25–30]. Laminin, along with type IV collagen and perlecan/HSPG2, normally is deposited in the basement membrane separating epithelial and stromal compartments in soft tissues and surrounding axons and Schwann cells in peripheral nerves [31–34]. Laminin also is found in territorial matrices surrounding cancer cells, where its aberrant expression is associated with changes in cell migratory behavior [35, 36]. The mechanical behavior of the basement membrane influences tissue development and cancer cell behavior [37, 38]; in fact, the invasive breaching of the basement membrane by cancer cells is the first step of metastasis [21, 39, 40].

**FIG. 1.**
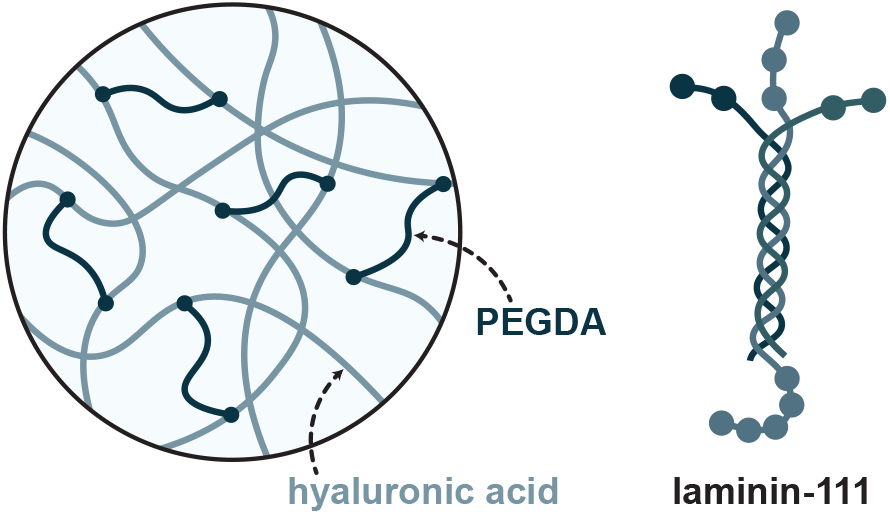
Molecular ingredients. Gels are constructed from thiol-modified hyaluronic acid polymers crosslinked by shorter poly(ethylene glycol) diacrylate (PEGDA) polymers. Varying concentrations of the glycoprotein laminin-111 are incorporated into the gels prior to crosslinking.

Interactions between cells and the composition and properties of their surrounding matrix directly determine cell behavior. In spite of the common use of ECM components in constructing three dimensional matrices and the recognition of the importance of large matrix deformations in cell and tissue morphogenesis, relatively little is known about the non-linear (large-strain) rheology of hydrogels of HA, laminin, or combinations of the two. Understanding the nonlinear mechanics of these gels is particularly important given that large deformations are inherently present in morphogenesis in native tissues and engineered ECMs. Given the unique multi-armed structure of laminin and its ability to polymerize into weak networks, it is conceivable that laminin could introduce unexpected mechanical properties into HA/laminin composite gels.

The modification of HA gel networks by inclusion of laminin has been shown to induce unanticipated changes in the differentiation behavior of organoid models [41], for which a mechanistic understanding is lacking. This is true in part because there remains a limited understanding of how laminin affects the nonlinear mechanics of hyaluronan-based hydrogels. To address this knowledge gap, we performed a series of rheological measurements on composite gels composed of HA, PEGDA, and varying concentrations of laminin. Using a parallel-plate rheometer, we probed the viscoelastic response of these gels under applied shear and axial deformation. Our primary goal is to determine how the inclusion of laminin-111 modifies the mechanical response of HA-PEGDA hydrogels, particularly in the nonlinear elastic regime under large strains.

## II. MATERIALS AND METHODS

### A. Gel preparation

HA-containing gels were prepared using HyStem® kits (GS1004F, Advanced BioMatrix) containing Glycosil® thiol-modified hyaluronan (MW 100 kDa) and degassed deionized water [42]. After HA reconstitution, Cultrex® 3-D Culture Matrix Laminin I (3446-005-01, BioTechne) was gently resuspended into the HA solution. Extralink-lite® poly(ethylene glycol) diacrylate (PEGDA) (MW 3500 Da) was then added and the combined solution was immediately pipetted into the rheometer. For laminin-only gels, the 6 mg/ml Cultrex® 3-D Culture Matrix Laminin I solution was pipetted directly into the rheometer.

All samples were allowed to polymerize for 90 minutes prior to rheological testing. The rheometer temperature was fixed at *T* = 37 ^°^C throughout gelation and all subsequent rheological tests.

### B. Rheological testing

Rheological measurements were performed using a TA Instruments DHR-3 Rheometer equipped with Peltier plate temperature control and an upper parallel-plate geometry of radius *R* = 1 cm at an initial gap separation *h* = 1 mm.

We employed a multi-stage shear rheology protocol, combining successive strain ramps with small-amplitude oscillatory shear testing at finite prestrain. A sample strain sequence is shown in Fig. S1. During the strain ramp stage, the shear strain *γ* (*t*) was increased at a constant shear rate 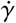 and the resulting stress *σ*(*t*) was measured. The differential shear modulus *K* is computed as

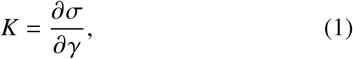

which in the small strain limit defines the linear shear modulus *G*_0_ = lim_*γ*→0_ *K*. All strain ramp data shown here correspond to the final ramp at shear rate 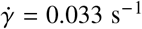 up to maximum strain *γ*_max_ = 2.0.

A finite-strain small-amplitude oscillatory shear protocol [43, 44] was applied immediately after each forward strain ramp to prestrain *γ*_0_, as shown in Fig. S1. For a series of increasing frequencies *ω*, we applied a superposed oscillatory shear strain increment *δγ* = 0.01, such that the time-dependent strain was *γ* (*t*) = *γ*_0_ + *δγ* sin (*ωt*). Fitting the resulting time-dependent shear stress *σ* (*t*) to *σ* (*t*) = *σ*_0_ (*γ*_0_)+ *δγ* [*K*′ (*γ*_0_, *ω*) sin (*ωt*)+ *K*″ (*γ*_0_, *ω*) cos (*ωt*)], we obtain fit values of the differential storage and loss moduli *K*′ (*γ*_0_, *ω*)and *K*″ (*γ*_0_, *ω*), respectively.

In the compression protocol, the axial strain *ε* (*t*) was decreased at a fixed strain rate 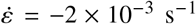 and the resulting normal stress *σ*_*N*_ (*t*) = *F*_*N*_ (*t*)/*A* was measured. Here, *F*_*N*_ is the normal force reported by the rheometer and *A* is the rheometer plate area. Note that negative strain values (*ε* < 0) correspond to compressive strain, and positive normal stress indicates a vertical normal force exerted by the gel on the upper plate of the rheometer. The strain-dependent apparent Young’s modulus *E*_app_ is computed as

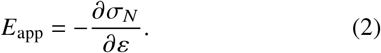

## III. RESULTS

### A. Response to applied shear

We first consider the influence of laminin concentration on the response of gels under applied shear. We primarily consider gels with an 8:1 mass ratio of HA to PEGDA (*c*_HA_ = 4.44 mg/ml and *c*_PEGDA_ = 0.55 mg/ml) and varying concentrations of laminin *c*_lam_. In Fig. 2, the differential shear modulus *K* is shown for these gels as a function of the shear stress *σ*, following the strain ramp protocol described in Section II B. We find that increasing the concentration of laminin leads to a moderate reduction in the linear shear modulus *G*_0_ (see inset of Fig. 2), while introducing some notable changes to the non-linear response. In the absence of laminin (*c*_lam_ = 0 mg/ml), we observe a smooth crossover from the linear regime to a stress-stiffening regime in which *K* ∝ *σ*^*a*^ with scaling exponent *a* ≈ 1. Such a linear scaling is consistent with recent theoretical predictions for fiber networks and experimental observations on reconstituted type I collagen networks [45]. Interestingly, a similar linear relationship between stiffness and stress is well-established for collagenous tissues under extension [46]. In Fig. S2, we show that similar stress-controlled stiffening behavior is seen both at other concentrations of HA and PEGDA and in gels composed of pure laminin. Notably, introducing laminin gives rise to a mild softening regime at intermediate strains, preceding the stiffening regime at larger strains. This softening regime becomes more pronounced with increasing *c*_lam_. We find that the stiffening exponent *a* appears to be insensitive to the laminin concentration, with *a* ≈ 1 for the concentrations we consider. However, the amount of stiffening in the large strain limit decreases with increasing laminin concentration. In all cases, the stiffening regime is followed ultimately by a failure regime, in which the apparent stiffness drops with increasing strain.

**FIG. 2.**
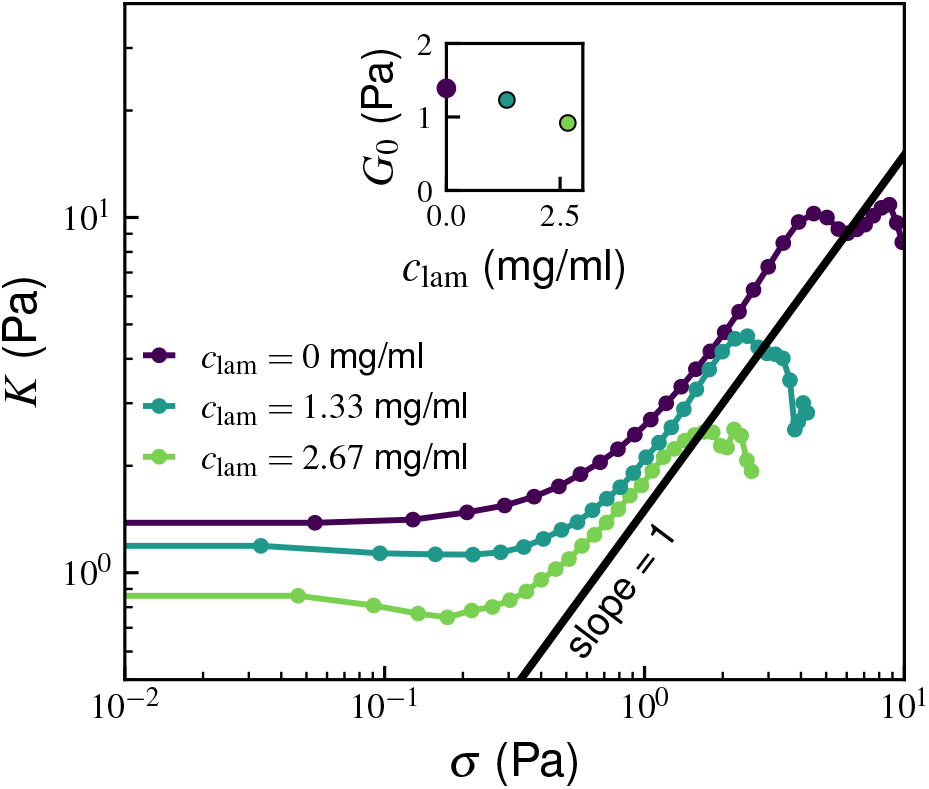
Stress-controlled stiffening under shear of crosslinked HA-PEGDA gels with varied laminin concentrations. Differential shear modulus *K* vs. shear stress *σ* for different concentrations of laminin *c*_lam_, following the strain ramp protocol. Here, the concentrations of HA and PEGDA are *c*_HA_ = 4.44 mg/ml and *c*_PEGDA_ = 0.55 mg/ml, respectively. The solid line indicates the scaling *K* ∝ *σ*^*a*^, with *a* = 1. Inset: The linear shear modulus *G*_0_ is plotted as a function of the laminin concentration *c*_lam_.

In Fig. S2b, we plot the differential storage modulus *K*′ (*γ*_0_, *ω*), measured using the prestrain-superposed small-amplitude oscillatory shear protocol described in Section II B, as a function of prestress *σ*_0_ for angular frequency *ω* = 0.21 rad/s. Here, we observe a similar stress-controlled stiffening regime with 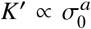, with *a* ≈ 1 for all samples. We also plot the loss tangent tan *δ* ≡ *K*^′′^/*K*′, which provides a measure of the dissipative character of the mechanical response (see inset of Fig. S2b). Notably, we find that increasing the concentration of laminin results in increased tan *δ*, suggesting that the incorporation of laminin leads to increased energy dissipation.

### B. Response to applied compression

We next consider the influence of laminin on the compressive response of our hydrogels. Fig. 3 displays the normal stress *σ*_*N*_ as a function of compressive strain −*ε* for the same gels as in Fig. 2, with an 8:1 mass ratio of HA to PEGDA (*c*_HA_ = 4.44 mg/ml and *c*_PEGDA_ = 0.55 mg/ml) and varying concentrations of laminin *c*_lam_. Increasing the concentration of laminin leads to minimal variation in the normal stress under small compression but decreases the normal stress considerably under larger strains, in comparison to the laminin-free gels. As was observed in gels under shear, the incorporation of laminin results in an intermediate softening regime preceding the stiffening regime. In the inset of Fig. 3, the apparent Young’s modulus *E*_app_ is plotted as a function of the normal stress *σ*_*N*_. The curves for varying laminin concentrations overlap reasonably well, all exhibiting a large-stress regime with

**FIG. 3.**
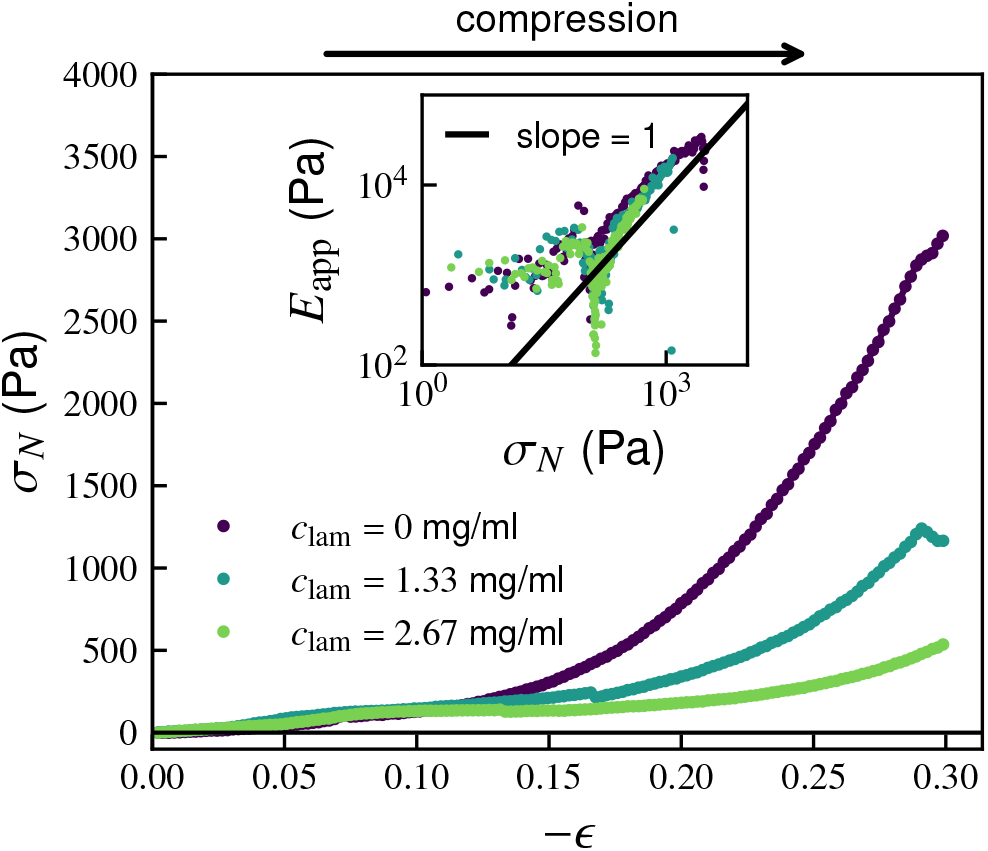
Nonlinear response in compression for crosslinked HA-PEGDA gels with varied laminin concentrations. Plot of normal stress *σ*_*N*_ vs. compressive uniaxial strain − *ε*. Inset: The apparent Young’s modulus *E*_app_ vs. normal stress *σ*_*N*_. The solid line corresponds to a slope of 1. Here, the concentrations of HA and PEGDA are *c*_HA_ = 4.44 mg/ml and *c*_PEGDA_ = 0.55 mg/ml, respectively.

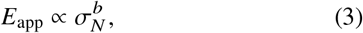

where stiffening exponent *b* ≈ 1. Similar to what we observed under shear, the stiffening exponent *b* appears to be insensitive to laminin concentration over the range of concentrations we consider.

## IV. DISCUSSION AND IMPLICATIONS

We have experimentally characterized the rheology of hydrogels composed of PEGDA-crosslinked HA and laminin under shear and compressive strains. We found that, for gels with fixed concentrations of hyaluronan and PEGDA, the incorporation of laminin leads to a reduction in the linear shear modulus *G*_0_ and changes the nonlinear mechanical response considerably. These findings suggest that laminin plays a nontrivial role in the nonlinear mechanics of composite gels, with potential implications for understanding cell-matrix interactions and morphogenesis in engineered ECMs.

Notably, while gels without laminin exhibit monotonic stiffening reminiscent of other crosslinked biopolymer hydrogels, the inclusion of laminin induces a mild strain-softening regime that precedes the eventual onset of strain stiffening. Similar intermediate strain softening behavior has also been observed in other gels [47, 48] and may be a consequence of the weak, noncovalent bonding and/or topology of the self-assembled laminin network. Consistent with this idea, we found that increasing the concentration of laminin resulted in an increased loss tangent tan *δ*.

Interestingly, in spite of the observed changes in the small- and intermediate-strain mechanical response due to the inclusion of laminin, we nonetheless find that in the large strain regime prior to failure, under both shear and compression, all gels exhibit a stress-controlled stiffness. Under shear, the differential modulus *K* scales as a power law with respect to the shear stress *σ* (*K* ∝ *σ*^*a*^ with *a* ≈ 1), consistent with behavior observed in other biopolymer networks such as reconstituted collagen gels [45]. Under large compression, the apparent Young’s modulus *E*_app_ scales as a power law with respect to the normal stress 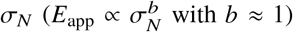.This stiffen-ing under compression is likely a consequence of the effective incompressibility of the gels; volume-preserving axial compression leads to lateral extension, such that the increase in the Young’s modulus *E* ultimately reflects the formation of tensile force chains. The observed exponents under both strain modes do not appear to depend on the concentrations of laminin. The robustness of this stress-controlled stiffening behavior under both shear and compression underscores its relevance for physiological and biomimetic ECM environments, in which cells and cellular collectives generically exert significant stresses on the surrounding matrix during self-assembly and regeneration.

Our results demonstrate that laminin acts as a strong modulator of the nonlinear mechanics of hyaluronan-based hydrogels. From a design standpoint, this reinforces the importance of accounting for nonlinear rheological behavior in engineered biomimetic ECMs for static and dynamic cell and tissue regenerative approaches [24, 49–56]. Matrix mechanics plays a major role in cellular self-assembly and organoid morphogenesis and has been studied extensively in various contexts, such as in salivary gland secretory structures and organoids [57–61]. The laminin-mediated intermediate strain-softening behavior that we observe could offer a mechanically permissive strain regime, possibly in tandem with biochemical cues, to facilitate shape change and reorganization in developing organoids [62, 63]. This could explain, in part, the observation in Ref. 41 that the inclusion of laminin in crosslinked HA gels is necessary for growth-factor-induced morphogenesis of salivary gland organoids [64].

Several open questions remain. The precise structure and distribution of the laminin within the HA-PEGDA network have yet to be elucidated. Future work could use fluorescence or electron microscopy to resolve whether the laminin forms, e.g., local aggregates or a well-distributed interpenetrating mesh with the HA-PEGDA network. A more complete picture of the structural features of the laminin component would provide crucial insight into the origins of the observed mechanical behavior.

Future experiments incorporating contractile cells within these matrices would enable direct measurement of how active forces interact with the observed nonlinear rheology. Previous studies of the rheology of cellularized fibrin networks have shown that contractile cells can actively stiffen their surrounding matrix [65]. It would be interesting to explore whether similar feedback effects occur in HA-laminin systems and whether the observed softening/stiffening transitions influence or are influenced by cellular activity and matrix deposition.

## ACKNOWLEDGMENTS

This work was supported in part by the National Science Foundation Division of Materials Research (Grant No. DMR-2224030), the National Science Foundation Center for Theoretical Biological Physics (Grant No. PHY-2019745), the National Institutes of Health (NIH/NIDCR R01DE022969, R01DE032364). JLS acknowledges support from the Eric and Wendy Schmidt AI in Science Fellowship, a Schmidt Sciences program.

## SUPPLEMENTARY INFORMATION

### S1. SHEAR RHEOLOGY PROTOCOL

All data shown here were gathered using a strain-controlled parallel plate rheometer. For shear tests, our strain protocol combined successive strain ramps of increasing magnitude with finite-strain small amplitude oscillatory shear tests of varying frequencies, in order to obtain estimates of both the differential modulus *K* (*γ*) and the differential storage modulus *K*′(*γ*_0_, *ω*). In Fig. S1, we plot a sample strain protocol. We found that, for some samples, the application of successive strain ramps led to creep, such that returning to the “zero strain” position yielded a negative shear stress. Although this is to be expected, it complicates our interpretation of *K* as a function of strain *γ*. To avoid this complication, we report *K* only as a function of the stress *σ* rather than strain *γ*.

**FIG. S1.**
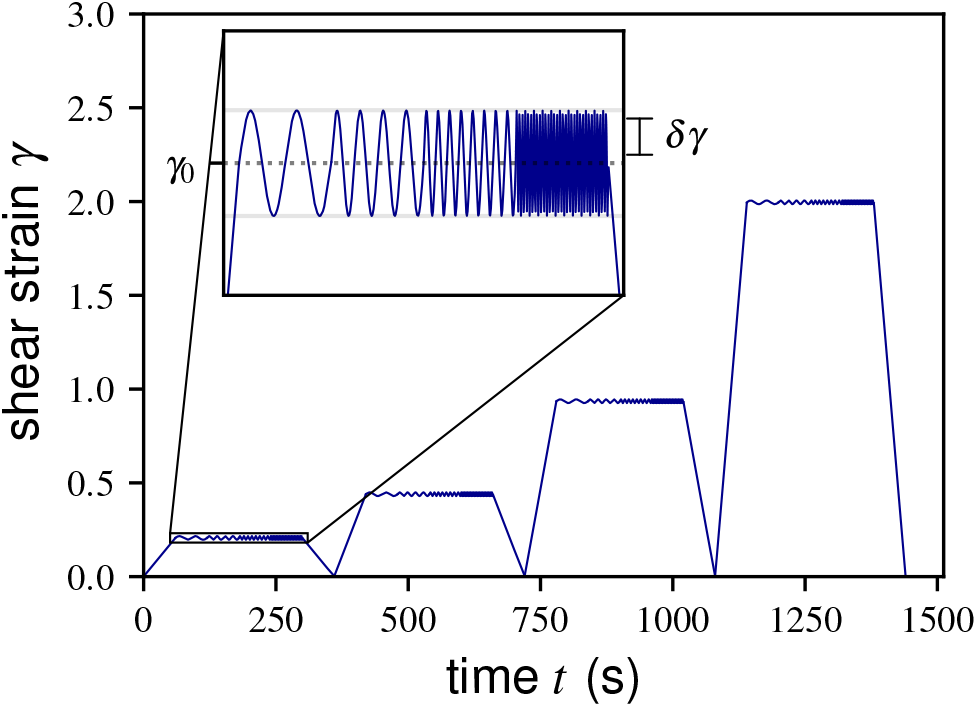
Multi-stage shear rheology protocol. The protocol combines successive strain ramps of increasing magnitude with small-amplitude oscillatory shear tests at varying frequencies. The applied shear strain *γ* is shown as a function of time *t* for a representative sample. The inset shows a magnified view of one of the oscillatory segments of amplitude *δγ* centered on prestrain *γ*_0_.

### S2. COMBINED STRAIN RAMP AND OSCILLATORY DATA

In Fig. S2a, we plot the measured differential modulus *K* as a function of stress *σ* for all samples under a strain ramp to maximum strain *γ*_max_ = 2.0. In Fig. S2b, we plot the differential storage modulus *K*′ and loss tangent tan *δ* ≡*K* ″ /*K*′ obtained via the oscillatory protocols.

**FIG. S2.**
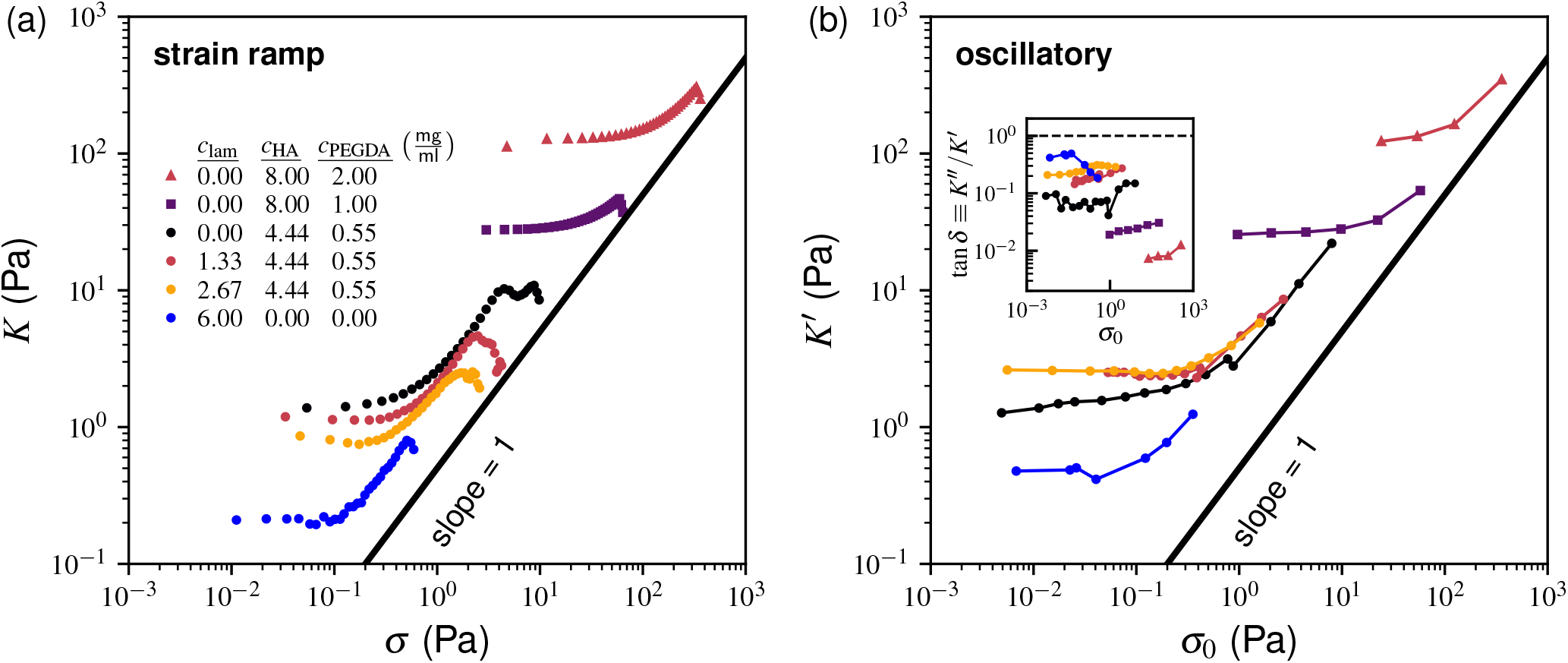
Stress-controlled stiffening across compositions. In panel (a), corresponding to systems undergoing the strain ramp protocol, the differential shear modulus *K* is plotted as a function of the shear stress *σ* for samples with varying concentrations of laminin, HA, and PEGDA, listed in the inset table. The solid line indicates the scaling *K*∝ *σ*^*a*^, with *a* = 1. Here, the strain rate is 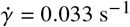 and the maximum strain is *γ*_max_ = 2.0. In panel (b), we plot the corresponding data for samples undergoing the oscillatory protocol: the differential storage modulus *K*′(*γ*_0_, *ω*) is shown as a function of the prestress *σ*_0_ (*γ*_0_). Here, *ω* = 0.21 rad/s. The inset shows the loss tangent tan *δ* ≡*K*″/*K*′ for the same data. The marker colors are identical to those in panel (a), and the lines connecting the markers are included only as a guide to the eye.

## Notes

### Competing Interest Statement

The authors have declared no competing interest.

